## supplemental file for "Distinct place cell dynamics in CA1 and CA3 encode experience in new environments"

### Materials and methods

#### Subjects:

All experimental and surgical procedures were in accordance with the University of Chicago Animal Care and Use Committee guidelines. For this study, 10-12 week old C57BL/6J wildtype (WT) male mice (23-33g) (4 WT for CA1 population imaging, Jackson Lab 000664) and C57BL/6-Tg(Grik4-cre)G32-4Stl/J (7 for CA3 population imaging, Jackson Lab, 006474) were individually housed in a reverse 12 hr light/dark cycle. Male mice were used over female mice due to the size and weight of the headplates (9.1 mm x 31.7 mm, ~2 g) which were difficult to firmly attach on smaller female skulls.  All training and experiments were conducted during the animal's dark cycle.

#### Mouse surgery and virus injection:

Mice were anesthetized (~1-2% isoflurane) and injected with 0.5 mL of saline (intraperitoneal injection) and ~0.45 mL of Meloxicam (1-2 mg/kg, subcutaneous injection). For CA1 population imaging, a small (~0.5-1.0 mm) craniotomy was made over the hippocampus CA1 (1.7 mm lateral, -2.3 mm caudal of Bregma). A genetically encoded calcium indicator, AAV1-CamKII-GCaMP6f (Addgene, #100834) was injected into CA1 (~75 nL) at a depth of 1.25 mm below the surface of the dura using a beveled glass micro pipette. For CA3 population imaging, the craniotomy was made over the CA3 (2.0 mm lateral, -1.7 mm caudal of Bregma). A Custom made Cre-dependent AAV virus: AAV1-CamKII-flex-GCaMP6f (made by Vigene) was injected (two injection sites at least 100 µm apart within the craniotomy, ~75 nl at each site) at a depth of 1.9 mm below the surface of the dura. After injection, the site was covered up using dental cement (Metabond, Parkell Coropration) and a metal head plate (Atlas Tool and Die Works). Water scheduling began the following day (0.8-1 mL per day and continued through all training and experiments). Around 7 days later, mice underwent another surgery to implant a hippocampal window as previously described[15]. Following implantation, the head-plate was reattached with the addition of a head-ring cemented on top of the head-plate which was used to house the microscope objective and block out ambient light. For CA3 mice, because the cannula window was implanted at an angle (~15 degrees) relative to the horizontal plane, we bent the two ends of the head plate to match this angle so that the head plate and cannula were on the same plane. We could then change the angle of our microscope objective to be perpendicular to this plane. Post-surgery mice were given 2-3 ml of water/day for 3 days to enhance recovery before returning to the reduced water schedule (0.8-1.0 ml/day).

Behavior and virtual reality (VR) switching:

VR and treadmill set-up were similar to previous studies[15,71]. VR environments (one training environment, which served as the familiar environment, F, and two novel environments: N1 and N2) were created using VIRMEn[72]. Each environment contained a 3-meter long linear track enriched with different distal and proximal 3D visual cues. 4 µL water rewards were delivered at the end of the track in all environments. During training, which began at least 5 days after window implantation, mice were placed in F for 30-40 mins each day and learned to run and lick the water reward in F. After each lap traversal, mice were teleported back to the beginning of the track. Before teleportation, a short VR pause of 1.5 s was implemented to allow for water consumption and to help distinguish laps from one another rather than them being continuous. Once mice reached the criterion > 2 laps per min that remained stable for 2-3 days (usually around 10-14 days after the start of training), imaging commenced.

Two-photon imaging:

Imaging was done using a laser scanning two-photon microscope (Neurolabware). The microscope consisted of an 8 KHz resonant scanning module (Thorlabs), a 16x/0.8 NA/3 mm WD water immersion objective (MRP07220, Nikon). GCaMP6f was excited at 920 nm with a femtosecond-pulsed two photon laser (Insight DS+Dual, Spectra-Physics) and the fluorescence was collected using a GaAsP PMT (H11706, Hamamatsu). The microscope is customized to tilt the objective, which we tilted to be perpendicular to the CA3 head plate angle but kept vertical for CA1 imaging. Stray light from the VR monitor was blocked from entering the objective lens by a dark rubber tube attached to the implanted head ring and the objective. Laser average power after the objective was ~60 mW for CA1 imaging and ~120 mW for CA3 to gain similar baseline florescence levels in the CA1 or CA3 FOV. Scanbox (Neurolabware) was used for microscope control and data acquisition. Time series videos were acquired at around 11 Hz for each of the 3 imaging planes (using an electronic lens) to maximize the number of neurons imaged in each mouse. The PicoScope Oscilloscope (PICO4824, Pico Technology) collected the signal from the microscope to synchronize frame acquisition timing with behavior (see below).

Imaging sessions:

Each mouse that reached the behavior criterion was carefully checked for expression under the two-photon microscope. Each mouse used in this study had healthy-looking GCaMP6f expression (resting fluorescence absent from the nucleus; fast transient kinetics; no signs of misshaped somas). On experimental day1: fields of view (FOV) were chosen that maximized the number of neurons across three planes. Imaging and behavior recordings started right before mice entered the VR. Mice ran at least 20 laps in F which took at least 10 min. After which the mice were instantaneously switched to a novel environment (N1). Mice then ran at least 35 laps in N1 and were recorded for at least 20 min and then placed back in their home cage. Experimental day 2: a similar procedure whereby mice were exposed to F first and then switched to a novel environment, but this novel environment (N2) was different from the first novel environment (N1). The FOVs were not necessarily the same as day1. After imaging, more than one averaged FOV was saved to be the reference for day 3 imaging in order to align the planes and record from the same cells the following day. Experimental day 3: The same FOVs were carefully matched to the previous day FOVs. Once imaging started, mice were directly exposed to N2 and ran for at least 29 laps and recorded for 20 min. Mice behavior including treadmill running speed, position, and licking were collected using the PicoScope Oscilloscope to synchronize with the imaging.

<u>Image processing and ROI selection:</u>

Time-series movies for multi-plane recordings were acquired using interleaved frames ($1^{st}$, $4^{th}$, $7^{th}$… frames belong to plane 1; $2^{nd}$, $5^{th}$, $8^{th}$… frames belong to plane 2: $3^{rd}$ ,$6^{th}$, $9^{th}$… frames belong to plane 3). Each multi-plane time series was then split into separate time series movies.

Same plane movies from Day1 in F and N1 were concatenated into one movie, as were Day 2 single plane movies in F and N2, and Day 2 N2 and Day 3 N2 single plane movies (for across days analysis of the same cells). Movement artifacts are corrected by customized MATLAB scripts based on whole frame cross correlation. For multiday imaging datasets (Day 2 N2 and Day 3 N2 concatenation), motion correction was applied before concatenation and then Fiji (ImageJ) was used to correct any rotational displacement between the two movies. The concatenated movies were then motion corrected again to assure the best alignment (Fig. 4).

Regions of interest (ROIs) were defined using customized MATLAB scripts as previously described[73] (mu = 0.6, 150 principal components, 150 independent components, s.d. threshold = 2.5, s.d. smoothing width = 1, area limits = manually chose for each FOV). For each ROI, baseline corrected ΔF/F traces across time, filtered for significant calcium transients were then generated as previously described[13,15,46].

Calcium transient analysis:

After extracting significant calcium transients, we analyzed and compared some basic characteristics of these transients across CA1 and CA3. Transient peaks: the maximum value for each transient from each neuron. Transient duration: the duration of each transient calculated at half peak from each neuron. Transient frequency: the frequency of significant transients from each neuron.

Behavior analysis:

First, immobile and backward moving periods were removed by identifying instantaneous velocity signals slower than 0.2 cm/s. Second, to calculate the mean lap velocity on each lap, we divided the track length (3 m) by the time taken to finish the lap. Third, to then calculate normalized mean lap velocity (Fig. 1b; Fig. 4b), we took the mean lap velocity on each lap and divided it by the mean velocity of the first 3 laps in F.

Defining Place fields:

Place fields were identified and defined as described previously[13,15,46]. Because mice ran continuously and consistently in all conditions, we included all laps and transients for place field identification. The 3 m track was divided into 50 bins (6 cm per bin). The mean ΔF/F was calculated as a function of virtual track position for 50 position bins for each lap, which formed a 50 by N laps matrix. Potential place fields were first identified as contiguous points of this matrix in which all of the points were greater than 15% of the difference between the peak ΔF/F value (from all 50 bins) and the baseline value (mean of the lowest 12 out of 50 ΔF/F values). The potential place field had to satisfy the following criteria to be defined as a significant place field: 1. The field width must be > 20 cm and < 150 cm. 2. The field must have at least one value bigger than 0.1 ΔF/F. 3. The mean in field ΔF/F value must be > 3 times the mean out of field ΔF/F value. 4. Significant calcium transients must be present on at least 15 laps out of all the laps that the mouse traversed. Potential place field regions that met these criteria were then defined as place fields if their P value from boot strapping was < 0.05, as described previously[46]. Place fields from cells that have multiple place fields used the same criteria and were treated independently. Transients that occurred outside of the defined place field region were removed for analysis of each specific field. The resultant place fields were then used in all subsequent analysis unless specified.

Histology and brain slices imaging:

We checked the CA3 expression of some of the Grik4-cre mice to ensure the GCaMP expression was restricted to CA3. Mice were anaesthetized with isoflurane and perfused with ~10 ml PBS followed by ~20 ml 4% paraformaldehyde in PBS. The brains were removed and immersed in 30% sucrose solution overnight before being sectioned at 50 µm-thickness on a cryostat. The brain slices were then collected on glass slides and mounted with a mounting media with DAPI (SouthernBiotech DAPI-Fluoromount-G Clear Mounting Media,010020). The whole brain slices were imaged under 10X with a Caliber I.D. RS-G4 Large Format Laser Scanning Confocal microscope from the Integrated Light Microscopy Core at the University of Chicago.

Spatial correlation:

To measure place field spatial correlation across environments, we found place cells that had place fields in either environment and then calculated the Pearson's correlation coefficient between the mean activity along the track (in 50 bins) for all laps in two environments. To measure the place field correlation within environment, we divided the session up onto two halves based on the total number of laps completed. We then calculated the mean activity along the track for each half and calculated the Pearson's correlation coefficient. For cells with multiple place fields, only the first place field on the track was included. To measure place field spatial correlation across days, we found place cells that had place fields in both days and then calculated the Pearson's correlation coefficient between the mean activity along the track (in 50 bins) for the last 10 laps in N day 1 and first 10 laps in N day 2.

Place field onset lap:

To determine place field onset lap (Fig. 2c-e, Fig. 4h), starting from lap 1 we searched lap-by-lap for a lap with a significant calcium transient present within the boundaries of the future place field calculated from all the laps in the session. Once the lap was found, we would then search for significant calcium transients on each of the next 5 laps. If 3 of the 6 laps had significant calcium transients within the place field boundaries, that would be considered the place field onset lap, if not, we move to the next lap and repeated the analysis. If we changed this criterion and instead used 2 out of 6 laps or 4 out of 6 laps to define place field onset lap, the differences in distributions we observed between CA1 and CA3 remained. To control for the different numbers of laps that the mouse ran in F and N, the comparison in Fig. 2d only included the first 25 laps in F and N. If we instead included the later laps in N, the result did not change.

Place field COM and spatial precision:

To calculate the spatial precision, we first calculated the somatic transient center of mass (COM) on each traversal along the linear track. We measured ΔF/F in each bin. We then used the following equation to calculate the COM for each traversal n ($COM_n$):

$$COM_n = \frac{\sum_i DF_i \cdot x_i}{\sum_i DF_i}$$

Where $DF_i$ is the somatic ΔF/F in bin i and $x_i$ is the distance of bin i from the start of the track. We then calculated the weighted average COM ($COM_w$) from all traversals n ($COM_n$ from each traversal was weighted by the peak transient ΔF/F on that traversal ($A_n$)):

$$COM_w = \frac{\sum_n A_n \cdot COM_n}{\sum_n A_n}$$

Spatial precision[13] (SP) was then calculated as follows (inverse of the COM standard deviation):

$$SP = \frac{1}{\sqrt{\frac{\sum_n A_n (COM_n - COM_w)^2}{\sum_n A_n}}}$$

Out/in place field firing ratio:

This was computed as the ratio between the mean ΔF/F in bins outside the place field and the mean ΔF/F in bins within the place field.

Position Decoding Analysis:

Based on a recent study[74], we chose to use the long short-term memory (LSTM) neural network model to test whether representations in CA1 were better at decoding position on the first lap of N than CA3 representations.

Considering the differences in number of place fields we measured in CA1 and CA3 (more PFs in CA1 than CA3), as well as the amount of data required to build a useful LSTM model, we grouped data from all CA1 and CA3 mice that ran more than 45 laps in N2, and matched the number of cells used from CA1 and CA3 to build the models.

To match the data from different mice and use the decoders to decode position on a lap-by-lap basis, we first changed the time-series based data to position based data: the 3 m track was divided into 100 bins (3 cm per bin). The mean ΔF/F was calculated as a function of virtual track position for 100 position bins for each lap. By doing this, data within and across mice became the same length by lap. CA1 data were then grouped into one dataset and CA3 data grouped into another dataset. The decoding data was restricted to periods when the animals were running.

To test decoding ability on the first lap in N, we first tested different parameters (LSTM[78,79] model network structure and the number of place cells used to build the models) to make sure that the two decoders had similar decoding ability in the later laps (validation set) (Supplementary Fig. 3). We chose a one-layer LSTM decoder with 1024 units to decode the animals' position from the input of 200 place cells. When building a model, 200 place cells were randomly chosen from the entire CA1 or CA3 place cell population, the data from the 6$^{th}$ to 35$^{th}$ laps were used to train the decoder to decode the animals' position on the track which had been divided to 50 bins. The 36$^{th}$ to 40$^{th}$ lap data were used as the validation set. The decoders were then used to decode the animals' position on the first lap based on the place cell activity on this lap. We repeated this procedure 20 times for CA1 and CA3 and compared the predicting error on the first lap between the regions. The predicting error was calculated as the mean of the absolute difference between the prediction position and the animals' real position on the first lap.

We also built a naïve Bayes decoder with the same data, though the decoding ability is not as good as the LSTM decoder for the validation set, we got the same result as the LSTM decoder for the first lap position decoding (that is the CA1 is better at decoding position on the first lap compared to CA3) (Supplementary Fig. 3).

Place Field Shifting:

To calculate population place field shifting (Fig 3e, 5d, 6c-d), we first calculate the COM for all place fields on each lap with a sliding window of 5 laps (the $COM_w$ of the current and 4 next laps). The 5-lap sliding average was done for smoothing purposes but the same trends were observed without smoothing. For each place field, lap-wise shift was computed as the difference between lap 12 and current lap. We could then calculate the average shifting over the population of place fields on each lap. Only place fields with place field activity on lap 12 and a place field onset lap < 20 were included. Also, due to the onset of place fields on different laps, the number of samples that contribute to the mean on each lap is different.

Note that we also used the 5-lap COM weighted average method in Fig. 3d.

Place Field Skewness

Place field skewness is calculated as the third statistical moment of the place field. For lap n:

$$Skewness_n = \sum_i \frac{DF_i}{\sum_i DF_i} \cdot \frac{(x_i - COM_n)^3}{\sigma^3}$$

where $\sigma$ is the place field's "standard deviation":

$$\sigma = \sqrt{\sum_i \frac{DF_i}{\sum_i DF_i} \cdot (x_i - COM_n)^2}$$

To calculate population place skewness trend, we first calculate the skewness for all place fields on each lap. Then we took each place field and aligned them together by the actual laps.

Place field width

Place field width on each lap is calculated as the difference between the first and last bin with in-field activity. We then normalized the lap-wise width to the mean PF width for each PF.

Notice, for COM shift and reset, PF skewness and width analyses, most PFs near start or end of the track were excluded if they were clipped, using the following criterion: if the distance between one track edge and PF COM was at least 1 bin shorter than the other half of the PF, the PF was excluded. For cells with multiple place fields, only the first PF on the track was included.

Statistical analysis

Error in the text and figures are presented as mean ± S.E.M, unless stated otherwise.

We used either an estimation approach or null-hypothesis testing to compare data (described in figure legends). To generate Gardner-Altman estimation plots, which highlight the effect size, we used the Data Analysis with Bootstrapped-coupled ESTimation (DABEST) package[74] (available on GitHub: https://github.com/ACCLAB/DABEST-python). To assess the uncertainty of the effect size, the mean difference between two distributions and its 95% confidence interval were bootstrapped. For null-hypothesis testing, Wilcoxon rank-sum test, Wilcoxon signed-rank test or Kolmogorov-Smirnov 2 sample test were applied. $P < 0.05$ was chosen to indicate statistical significance and $P$-values in figures are indicated as follows: ∗, $P < 0.05$, ∗∗, $P < 0.01$, ∗∗∗, $P < 0.001$, N.S. not significant. For data tested with the estimation approach, we also used the null-hypothesis testing to confirm any differences.

For linear regressions of the population slopes (e.g. Fig. 3e left), the $P$-value for the t-statistics to test whether the slope was significantly positive or negative were reported following the same approach reported above. To compare the population dynamics of different conditions, we performed exact testing based on Monte-Carlo resampling[77] (1000 re-samples with sample size matching the lower sample size condition) as detailed in legends.

To assess the shifting dynamics in single PFs, we performed linear regression on the lap-wise $COM_n$ relative to onset lap (Fig. 3c), and significance was assessed with an F-test.

<u>Data and Software:</u>

Data processing and analysis was performed using custom written scripts in MATLAB or Python.

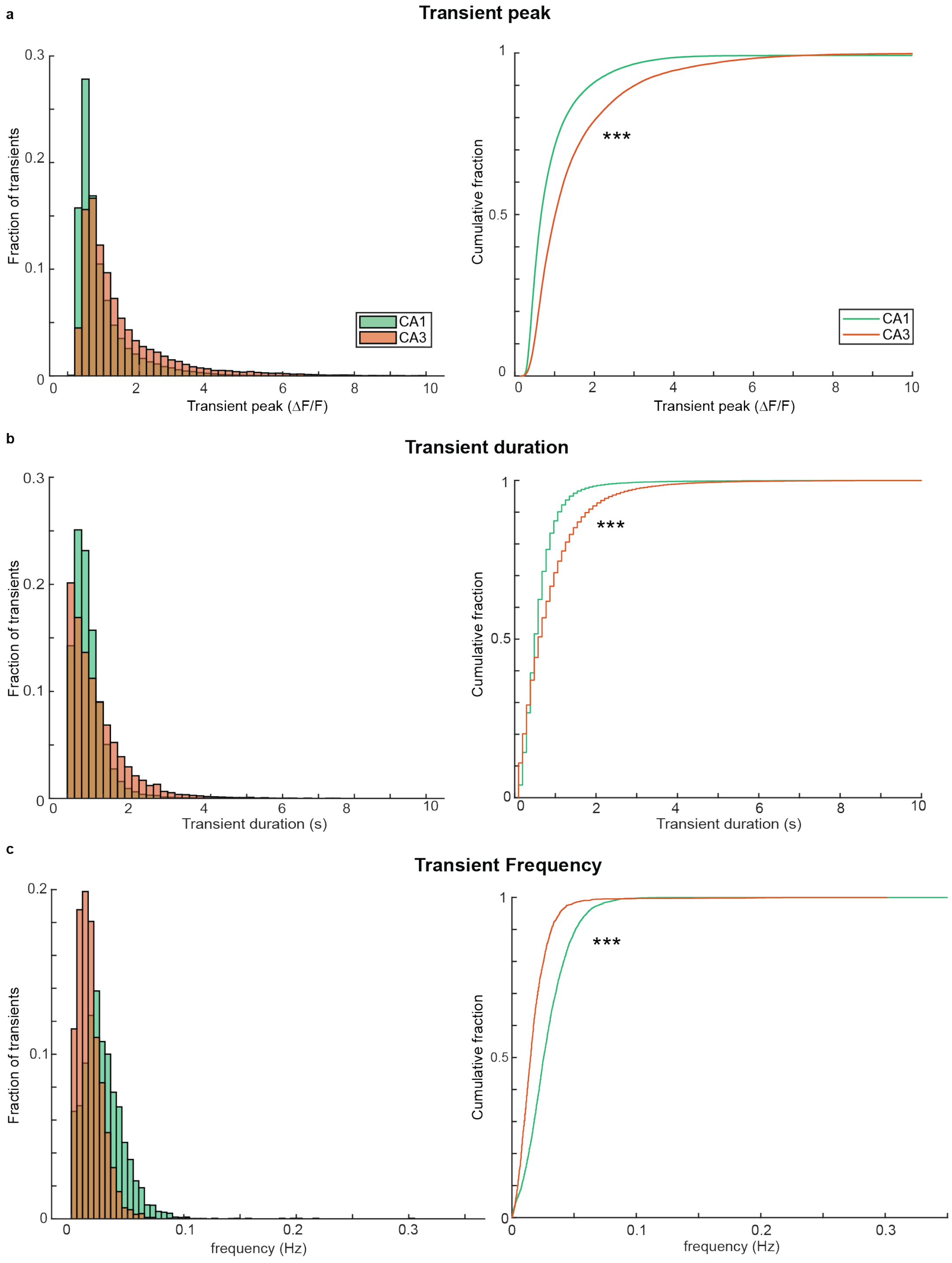


**Supplementary Fig. 1 Calcium transient properties in CA1 and CA3.**

**a,** Left, histogram of the Calcium transient peaks for CA1 (green) and CA3 (orange). Right, the corresponding cumulative fraction plot. Wilcoxon rank sum test, ***, $P < 0.001$. **b,** Left, histogram of the Calcium transient durations calculated at 50% of the peak. Right, the corresponding cumulative fraction plot. Wilcoxon rank sum test, ***, $P < 0.001$. **c,** Left, histogram of Calcium transient frequencies. Right, the corresponding cumulative fraction plot. Wilcoxon rank sum test, ***, $P < 0.001$.

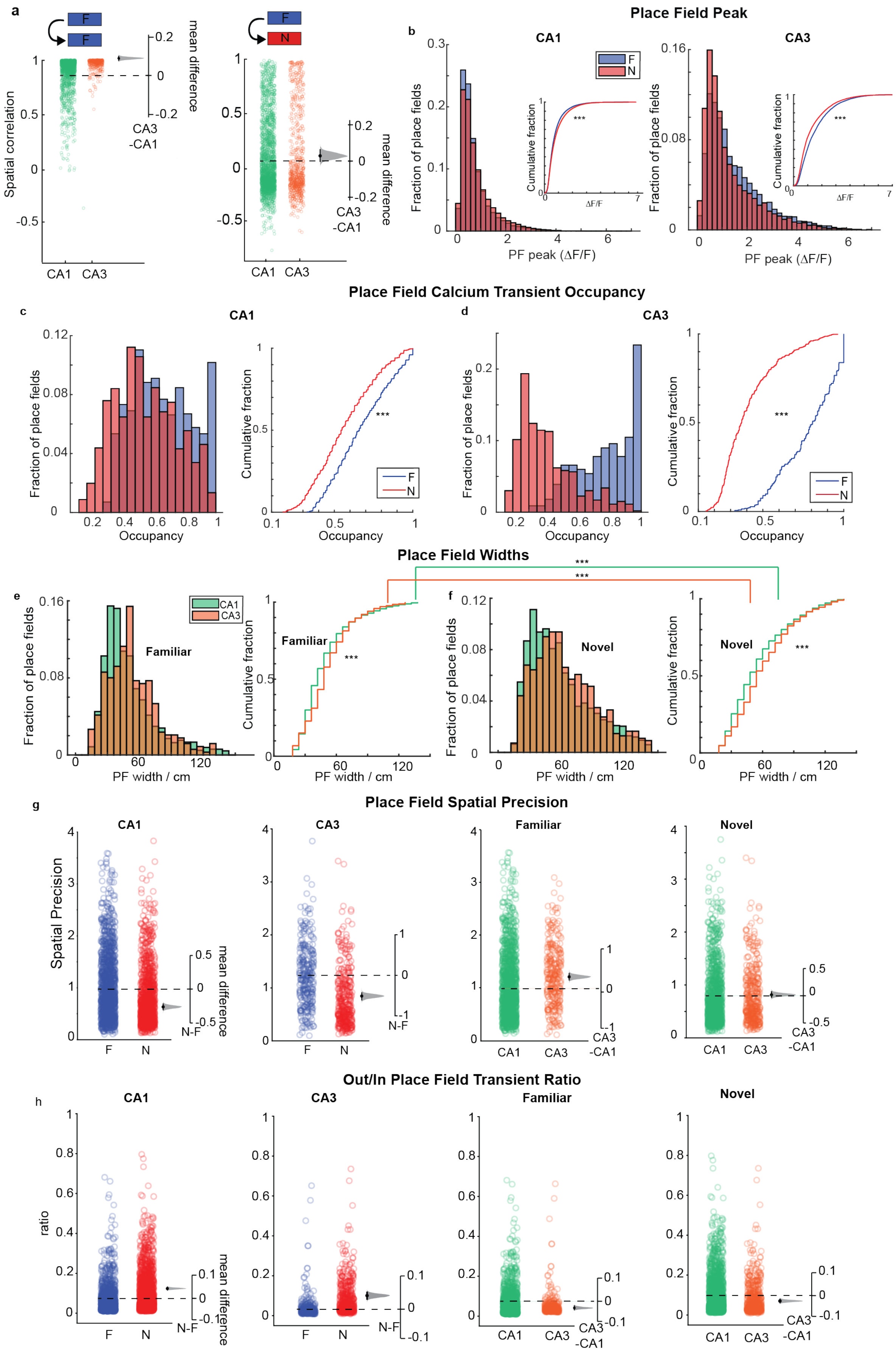

a
F
F
N
Spatial correlation
mean difference
CA3 -CA1
CA1 CA3
b
Place Field Peak
CA1
CA3
Fraction of place fields
PF peak (ΔF/F)
Cumulative fraction
ΔF/F
***
c
Place Field Calcium Transient Occupancy
d
Occupancy
e
Place Field Widths
Familiar
Novel
CA1
CA3
PF width / cm
f
g
Place Field Spatial Precision
Spatial Precision
N-F
h
Out/In Place Field Transient Ratio
ratio

**Supplementary Fig. 2 Place field properties in familiar and novel environments in CA1 and CA3.**

**a,** Pearson's correlation coefficient of each cell's mean place field (PF) between CA1 and CA3 within familiar (F) sessions (left) and between F and novel (N) sessions (right). Bootstrapped mean difference (Δ) shown on the right of each plot. Note CA3 PFs are much more stable within F than CA1 PFs. **b,** Histogram of each PFs peak activity for CA1 (left) and CA3 (right) in F (red) and N (blue). Insets show cumulative fraction plots. Wilcoxon rank sum test, ***, $P < 0.001$. **c,** Left, histogram of PF Calcium transient occupancy (percentage of laps with PF activity) in F (red) and N (blue) in CA1. Right, corresponding cumulative fraction plot. Wilcoxon rank sum test, ***, $P < 0.001$. **d,** Same plots as (c) in CA3. Note the striking difference in transient occupancy in N versus F in CA3. The lower transient occupancy in N is largely due to delayed PF formation in N in CA3 (See Fig. 2). **e,** Histogram of each PFs width in CA1 (green) and CA3 (orange) in the familiar environment. Right, corresponding cumulative fraction plot. Wilcoxon rank sum test, ***, $P < 0.001$. **f,** Same plots as (e) but in N. **g,** Spatial precision of each PF in CA1 (left) and CA3 (middle left) in F (blue) and N (red). Same data are also compared across CA1 and CA3. Bootstrapped mean difference (Δ) shown on the right of each plot. Note that PFs in N are less precise than in F in both CA1 and CA3, and CA3 PFs in F are more precise than CA1 PFs in F. **h,** Ratio of out versus in PF activity for CA1 (left) and CA3 (middle left) in F (blue) and N (red). Same data are also compared in F (middle right) and N (right) across CA1 and CA3. Bootstrapped mean difference (Δ) shown on the right of each plot. Note that PFs in N have more out-of-field firing than F in both regions but CA3 has less out-of-field firing than CA1.

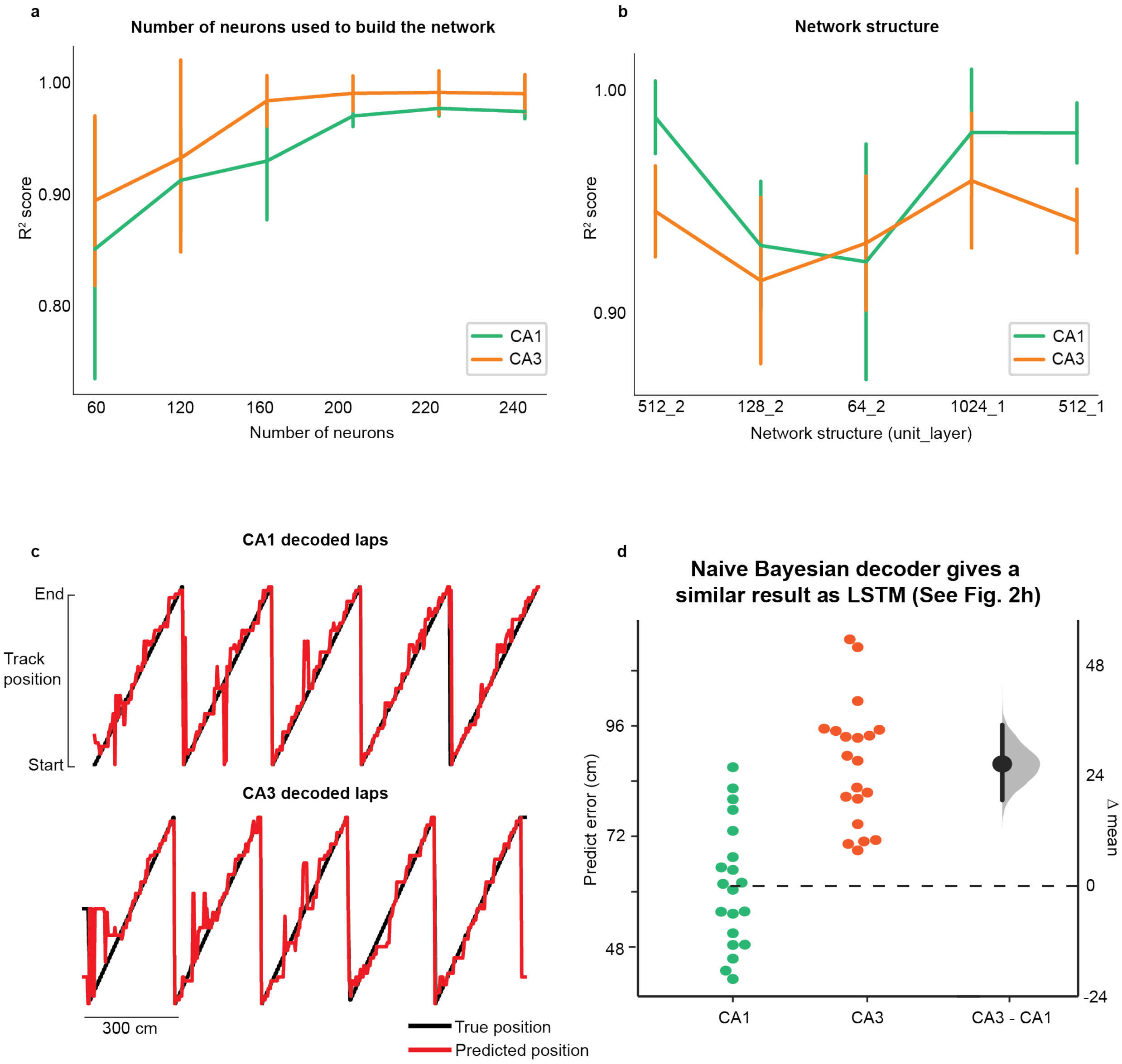


**Supplementary Fig. 3 Long-Short-Term-Memory (LSTM) and Bayesian decoding of animal position.**

**a,** Mean LSTM decoder performance ± SEM versus the number of place cells that were used to build the decoder. The network structure for all the decoders are the same, a one-layer 1024-unit neuronal network. **b,** Mean decoding performance ± SEM versus the network structure for the decoder. All decoders were trained with 200 place cells. **c,** Example mouse showing true track position (black) on laps 36-40 and the predicted position (red) decoded by a Naïve

Bayesian decoder. Note the LSTM decoder (Fig. 2f) did a much better job than the Bayesian decoder. **d,** Average predicted error for the first lap in the novel environment for CA1 and CA3. Each dot represents the decoding error for one decoder trial built based on the activity of 200 randomly chosen place cells from CA1 (green) or CA3 (orange) data. *n* = 20 decoder trails. Bootstrapped mean difference (Δ) shown on the right of each plot.

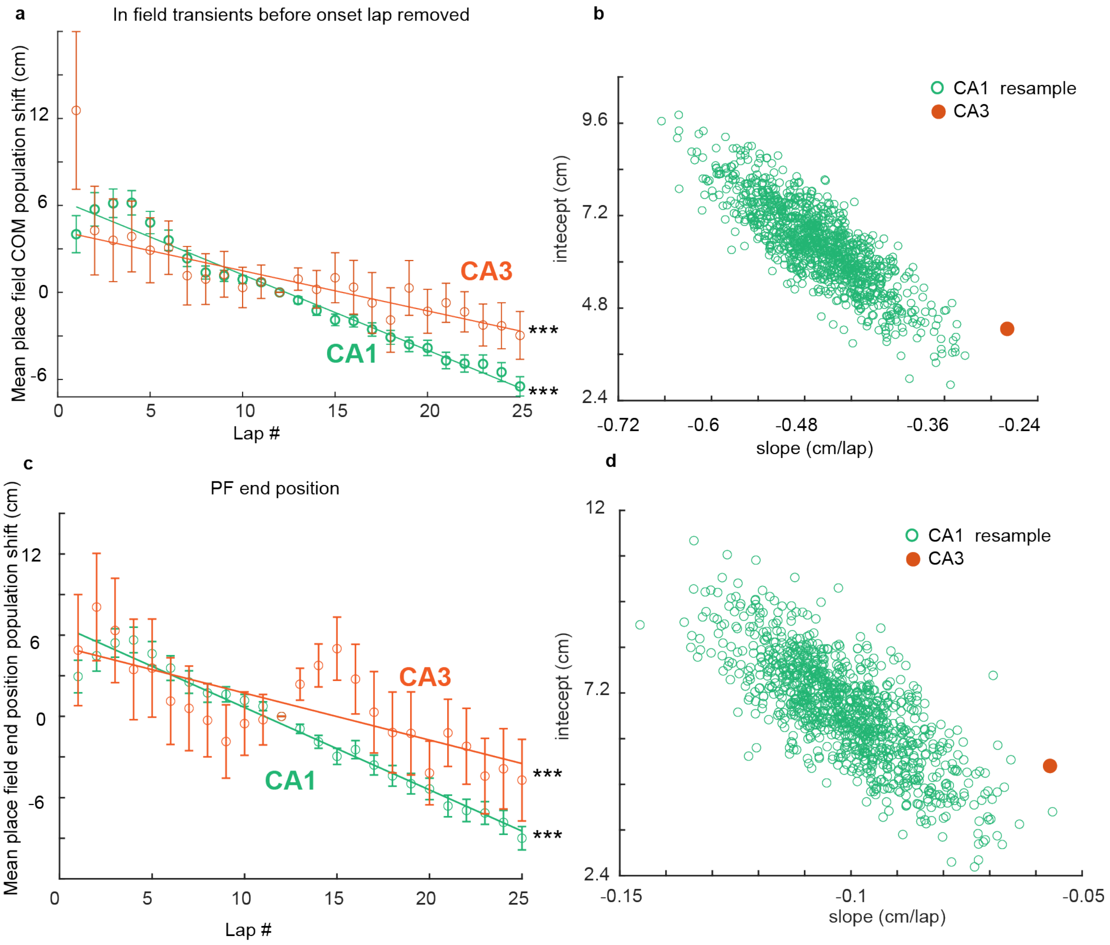


**Supplementary Fig. 4 Population backward shifting is still observed when transients that occur prior to place field emergence are removed and when place field position on each lap is defined by the location of the end of calcium transient.**

**a,** Same as Fig. 3e but with all in-field transients before each place field (PF) onset lap removed. ***, $P < 0.001$. **b**, Resampling analysis (1000 resamples) associated to (a) shows CA1 backward shifting is still significantly faster than in CA3. **c**, Same as Fig. 3e and (a) but with PF position on each lap not defined by the COM but by the location of the end of the fluorescence transient. This shows that the location of the end of the PF shifts backward in both CA1 and CA3. **d**, same as (b) but associated to (c): CA1 population shifting slopes are still significantly more negative than CA3 (*P* = 0.001).

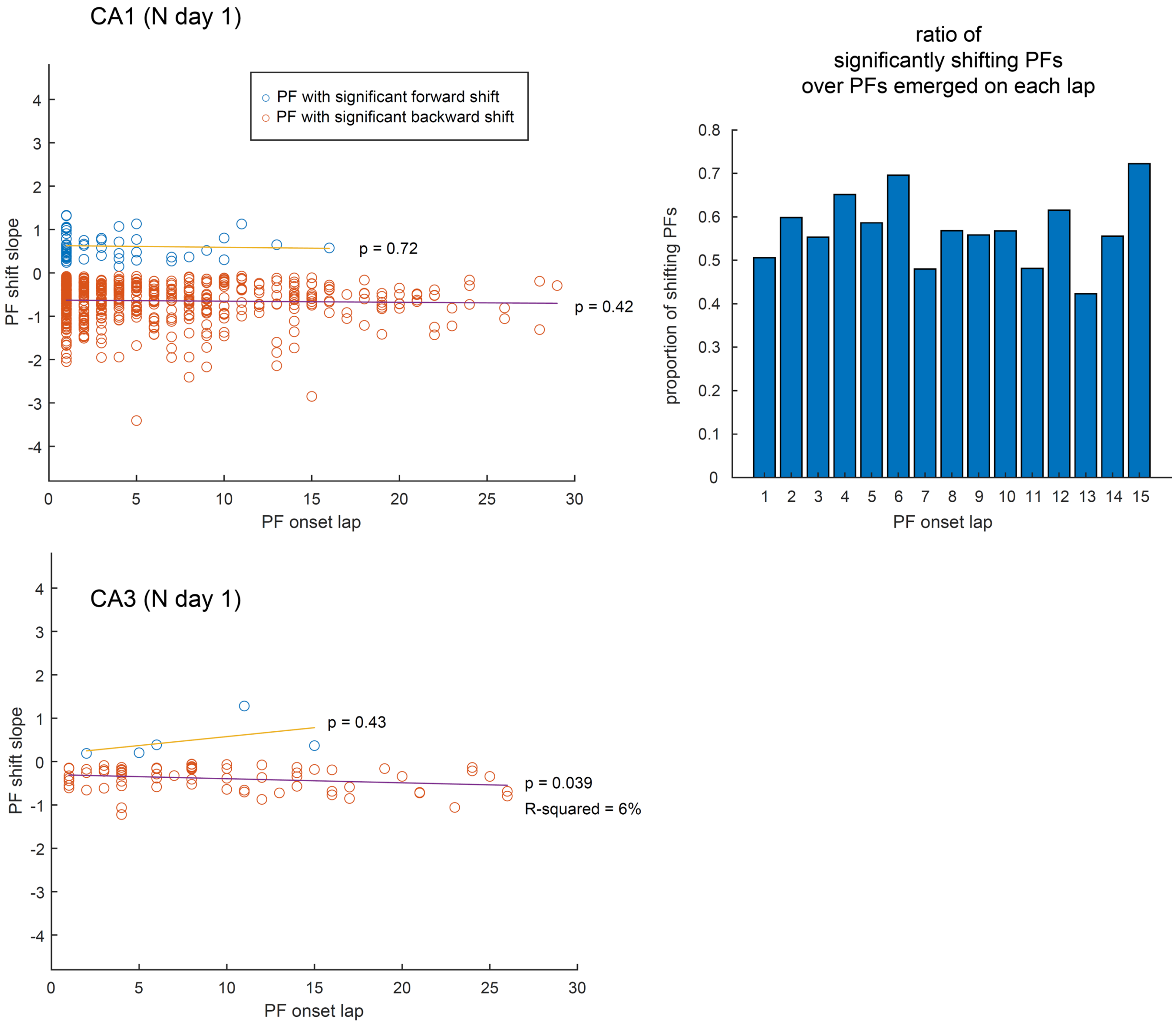


**Supplementary Fig. 5 Lap-by-lap shifting is not related to the timing of PF emergence**

Place field (PF) shift slope is defined as in Fig. 3c. Only significantly shifting PFs (regression $P < 0.05$) are included. No correlation between PF onset lap and the amplitude of their shifting dynamics is observed in CA1, and only weakly in CA3 where the sample size is low. Note that, in CA1, the higher number of shifting PFs with early onset is simply due to the higher number of PFs emerging in early laps. The proportion of shifting PFs (Right panel) stays stable throughout the first 15 laps (later laps not shown because number of shifting PFs < 10).

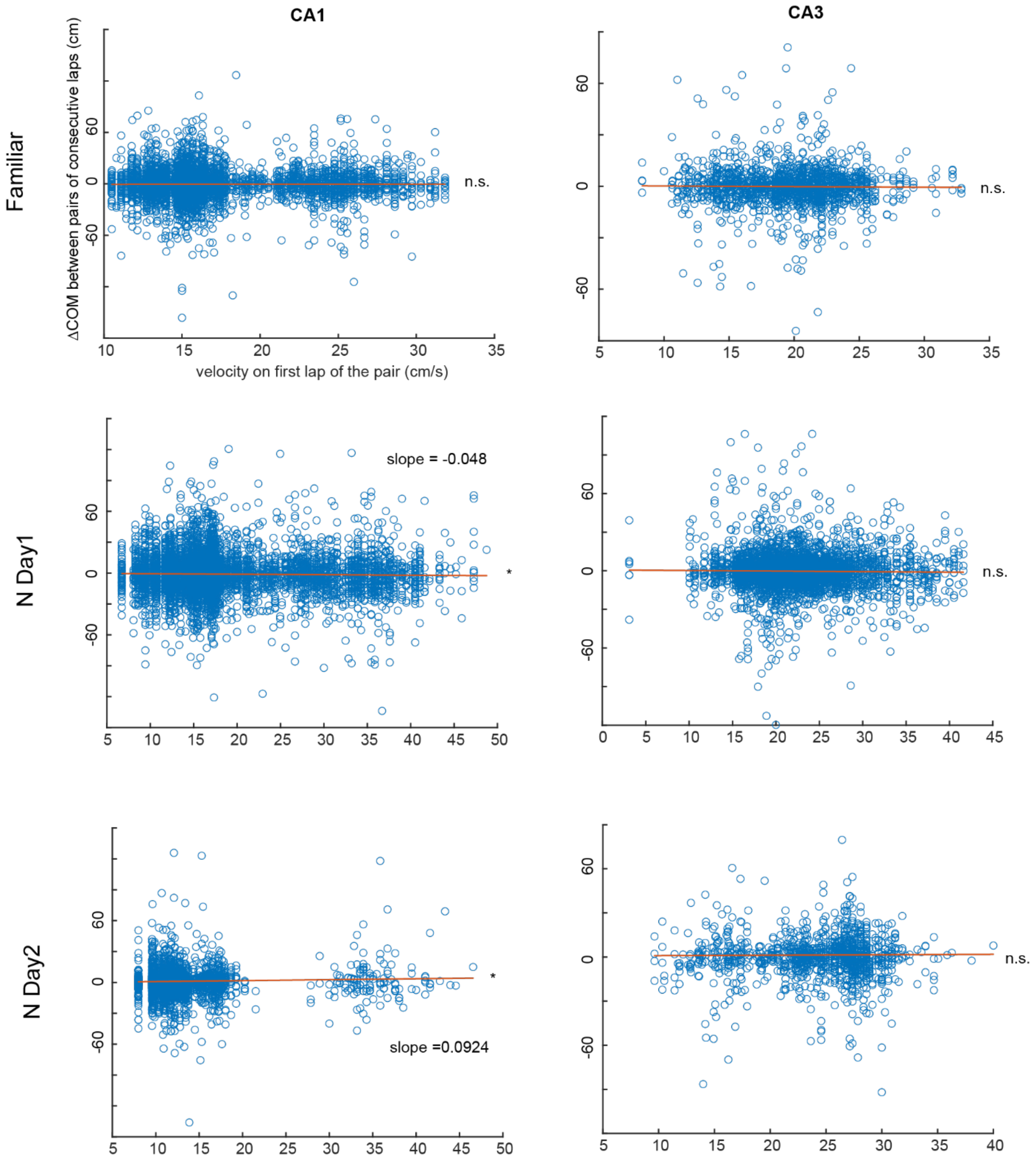


**Supplementary Fig. 6 Relationship between place field shifting and lap velocity.**

For all place fields (PFs), all differences in COM position between consecutive laps (ΔCOM) are plotted against the velocity on the first lap of the pair (similar results when velocity on second lap is used instead). This analysis does not reveal an obvious relationship between velocity and shifting and clearly shows that large lap-to-lap shifts are not due to higher velocities.

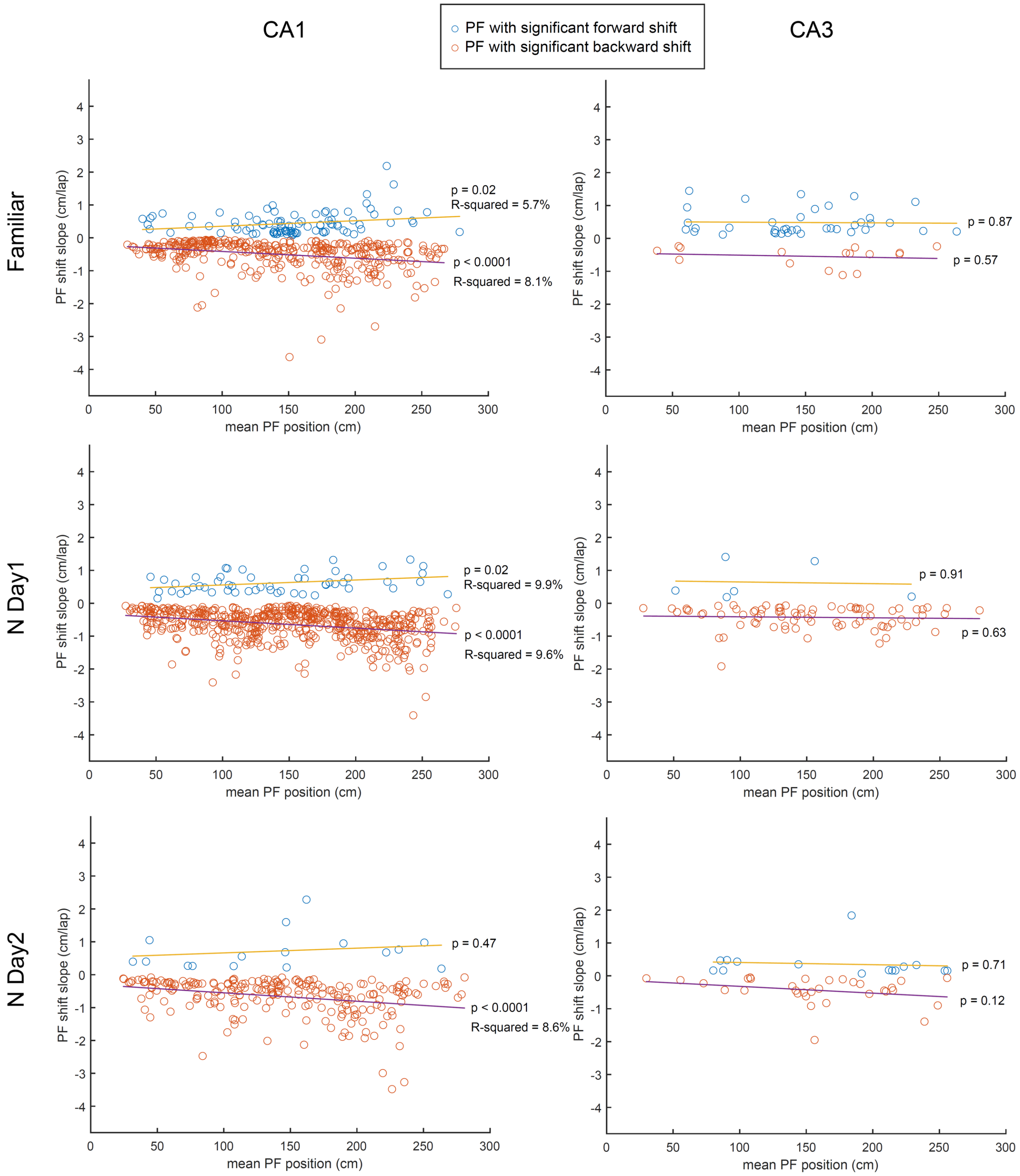


**Supplementary Fig. 7 Place field shifting is weakly correlated to place field position.**

Each data point corresponds to the shifting slope of a single place field (PF) from the regression analysis on the PF COM lap-by-lap position (see Fig. 3c). Only PFs with regression *P*-value < 0.05

are included. In CA1, PF shifting amplitude is weakly but significantly correlated with the PF mean COM position in each recording day and environment. PFs with a large forward or backward shift are biased toward the end of the track. No correlation is detected in CA3 but the number of shifting PFs is much lower.

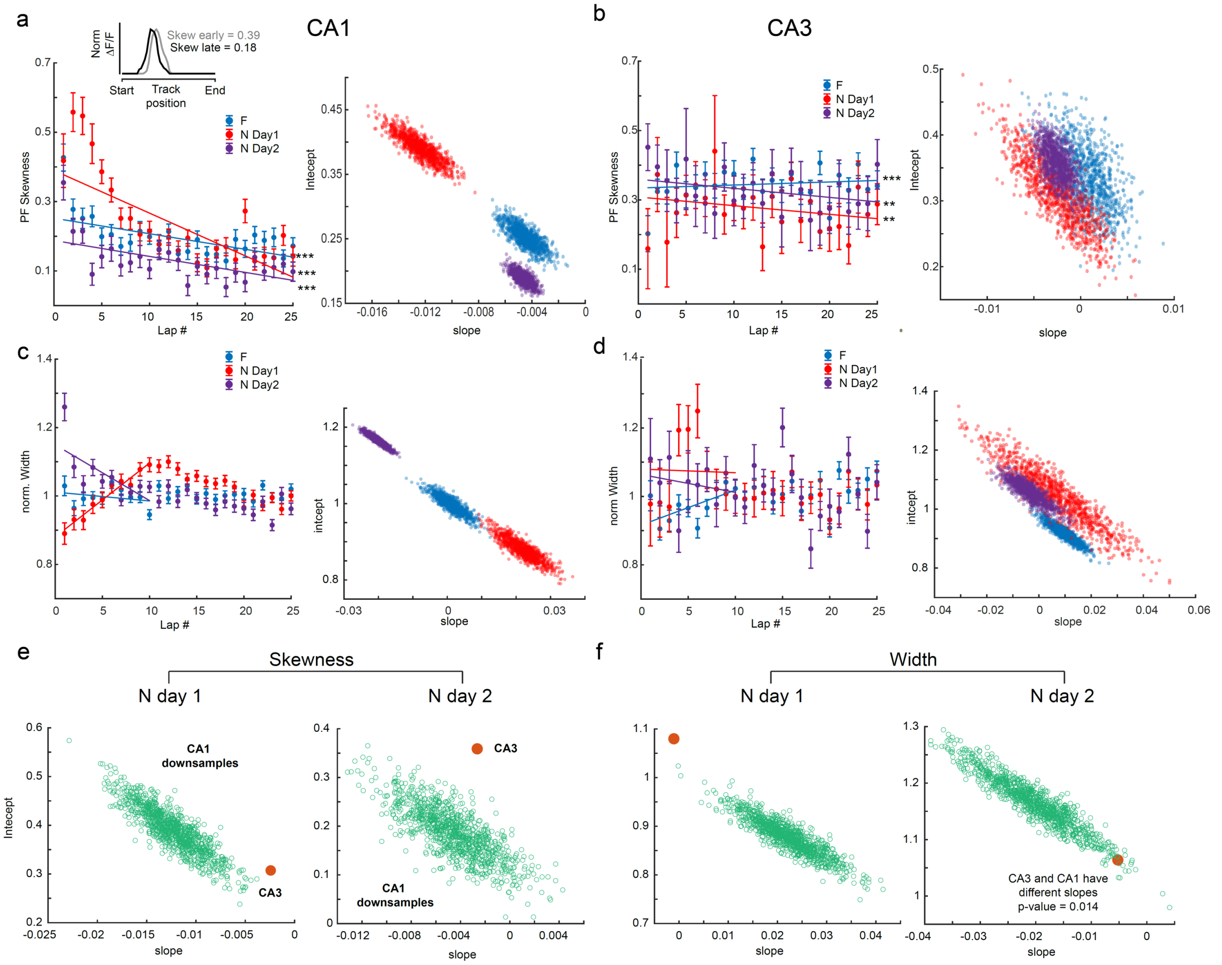


**Supplementary Fig. 8 Lap-by-lap change of place field skewness and width.**

**a,** Left, Lap-wise mean ± SEM skewness over all place fields (PFs) recorded in a given condition. Linear regression (on data points, not means) shows a decrease in positive skewness with experience, especially in N day 1. Inset: representative example PF that shifted backward, became less positively skewed and wider (early = lap 1-5 average fluorescence activity, late = lap 25-30 average). Right, resampling analysis as in Fig. 6c: slopes are significantly different from N day 1 to N day 2 and stay similar in a very familiar environment (F). Overall, these results suggests that PFs become less skewed over the first 10 laps in N and seem to stabilize afterwards, with slow population change upon re-exposure. **b,** same as (a) for CA3. Distribution spread is wide, suggesting less homogeneity in the population of PFs. Overlap in distributions show dynamics do not change significantly with familiarization. **c-d,** Same as (a) for the lap-wise PF width normalized to width averaged over all laps for each PF (i.e. 1 means that the width is the same as the average PF width). This normalization controls for large variations in individual PF width and allows direct comparison with Mehta et al. 1997[45]. In CA1, PF width increases over first 10 laps in N but decreases on second day and is stable in F. CA3 does not show clear population dynamics, although PFs become more homogeneous with familiarization across days. Linear regression fit to data points from the first 10 laps only. **e-f,** Resampling analysis as described in Fig. 3e. CA3 and CA1 PF width and skewness dynamics (slopes) are significantly different on day 1, but only widths are different on day 2.

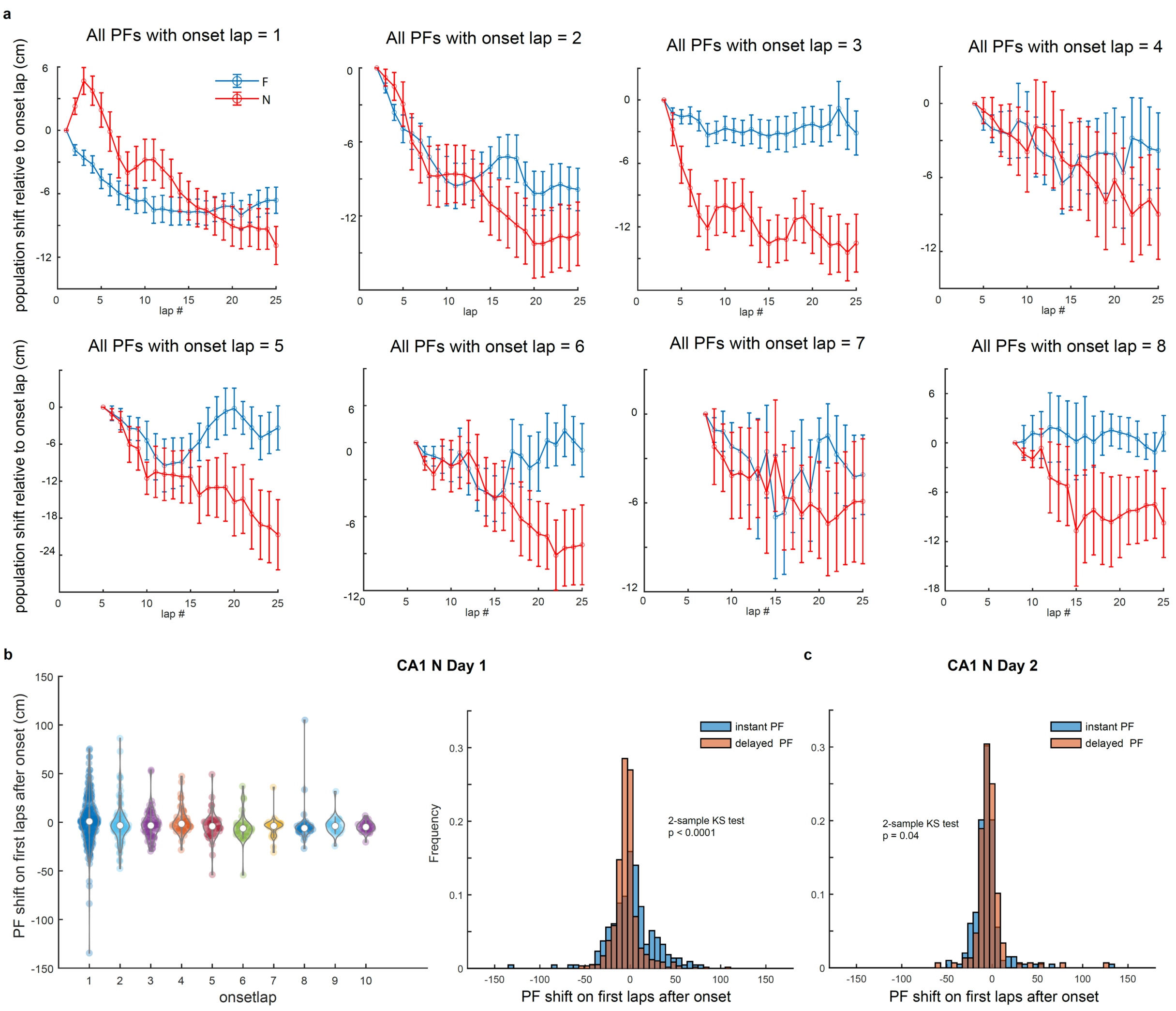


**Supplementary Fig. 9 CA1 forward shifting during first laps is driven by instant-onset place fields.**

**a,** Mean ± SEM, for all place fields (PFs), of 5-lap rolling average center of mass (COM) difference relative to onset lap, in novel (N) and familiar (F) environments. Forward population shifting during first laps after onset is only seen for instant PFs (onset lap = 1). **b,** Left, difference between 5-lap rolling average COM at onset lap and next lap, for each PFs, as a function of onset lap (not showing onset lap >10). The median is above 0 for instant PFs, not the other groups. Right, same as b but combining all delayed PFs together for comparison with instant

PFs. Distributions are significantly different, with more early forward shifting PFs in the instant PFs group. **c,** The effect seen in b disappears on day 2. Distributions are only mildly different and not because of a higher number of early forward shifting PFs in the instant onset group.

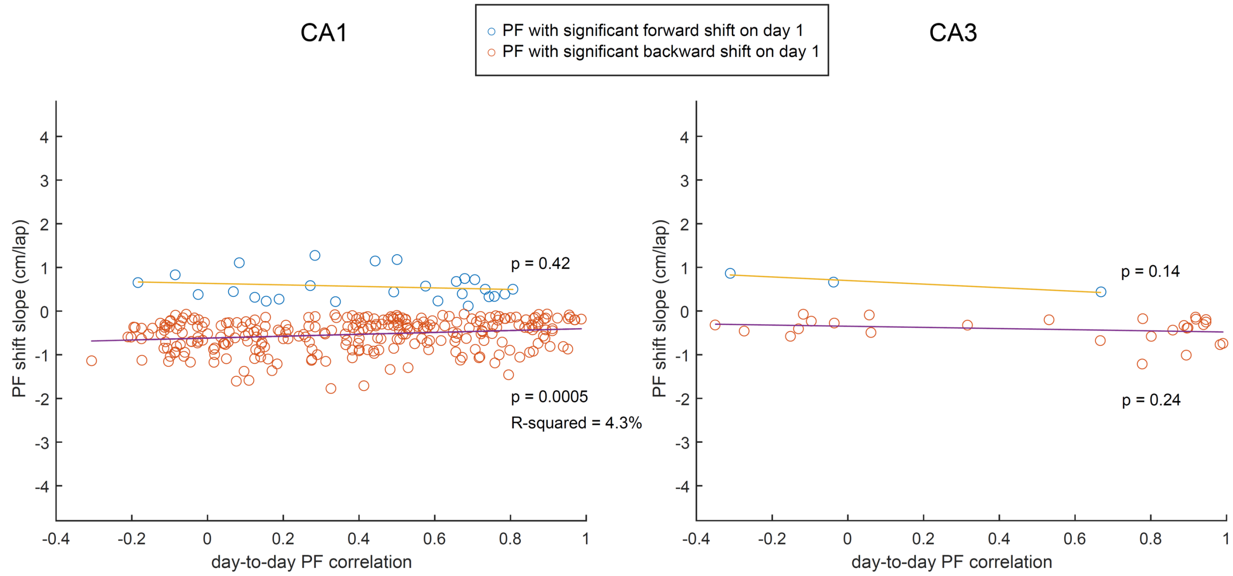


**Supplementary Fig. 10 CA1 place field stability across days is correlated to lap-by-lap stability on day 1.**

Place field (PF) shift slope is defined as in Fig. 3c. Only significantly shifting PFs (regression $P < 0.05$) are included. In CA1, there is a weak correlation (linear regression) between day-to-day stability and day 1 lap-by-lap dynamics for backward shifting PFs, suggesting that PFs that are stable across days are also more stable from lap-to-lap on day 1. Although not significant, this trend is seen for forward shifting PFs too. In CA3, PFs do not shift as much (see Fig. 3), resulting in a low sample size and no apparent correlation.

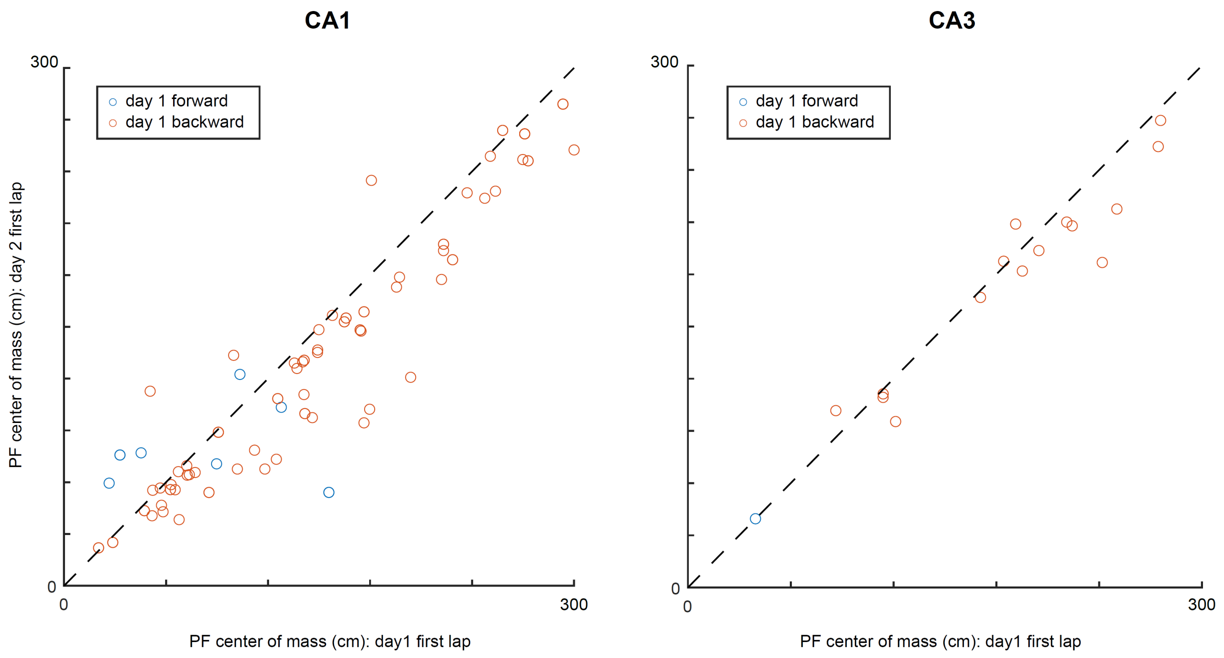


**Supplementary Fig. 11 On day 2, shifting place fields do not necessarily reset to their exact initial position on day 1.**

Place field (PF) shift slope is defined as in Fig. 3c. Selected PFs (day-to-day correlation > 0.5 and significant shifting on day 1) are the same as in Fig. 5b.

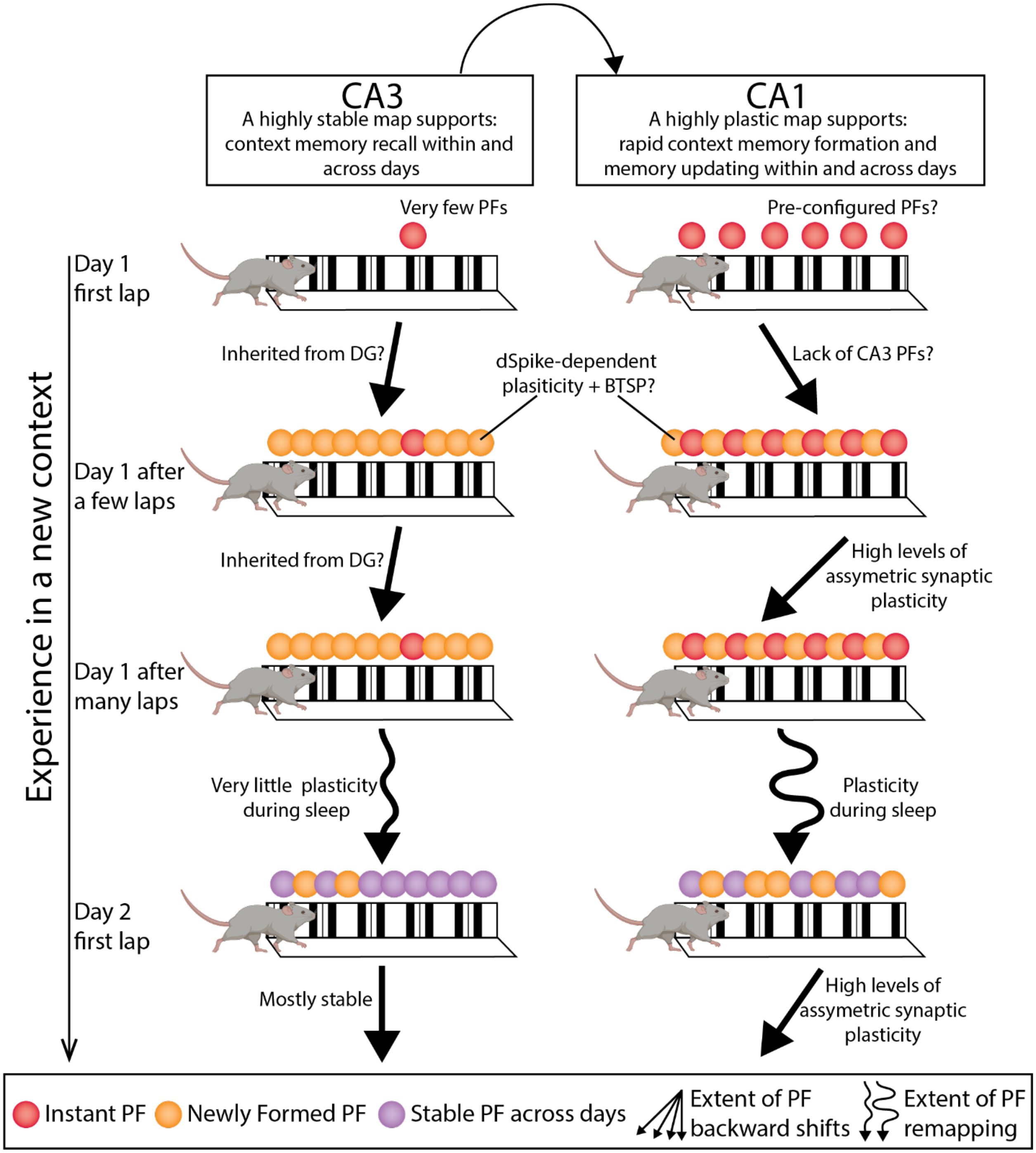


**Supplementary Fig. 12 Conceptual model.**

In a new context, place fields (PFs) in CA1 instantly appear on the first traversal (red circles).
These PFs do not require experience-dependent synaptic plasticity and so are likely pre-

configured[76]. Instant PFs are lacking in CA3. Instant PFs in CA1 initially shift forward for the first 1-4 laps, possibly due to the lack of CA3 PFs. Within the first few laps in the new context, new PFs appear in both CA1 and CA3 (orange circles). We partially know how these PFs form in CA1, but it remains unknown whether the same mechanisms are at play in CA3. In CA1, these PFs form through local dendritic spikes (dSpikes) that induce synaptic potentiation through an NMDA receptor mechanism[15]. This process occurs during the silent period when these cells are not firing. On the lap where the PF first appears, in some or many of these cells, behavioral timescale synaptic plasticity (BTSP) further strengthens synapses more globally throughout the neuron through burst firing associated with plateau potentials generated in the dendrites by coincident input from CA3 and Entorhinal cortex[24]. Throughout experience the newly formed PFs shift backwards and develop NMDA-dependent negative skewness in CA1 likely through asymmetric synaptic plasticity at the CA3-CA1 synapse. CA3 PFs show only a small amount of backward shifting and no increase in negative skewness, suggesting they may inherit their shifts from their inputs. Across days, PFs in CA1 undergo partial remapping, which likely involves synaptic plasticity that occurs during sleep when place cell sequences are known to be reactivated. CA3 PFs are much more stable across days. Throughout experience on day 2, a reduced but still very significant level of backward shifting occurs in CA1 with even less backward shifting in CA3. These PF dynamics in CA1 and CA3 likely support distinct roles in memory processing. We suggest that the CA1 rapidly forms a memory of a new context that supports single trial learning, and continuously updates this memory throughout ongoing experience. This updating - in the form of backward shifting PFs - may enable the CA1 to predict the near future regarding where in the context the animal is about to visit. The partial remapping across days in CA1 may also be another form memory updating by separating memory representations of experiences that occur in the same context on different days or at different times. On the other hand, animals need to recognize where they are and recall whether they are in a context they have experienced before. CA3 PFs seem to support this memory process by maintaining stable PFs within and across days.
